# IBD analysis of Australian amyotrophic lateral sclerosis *SOD1*-mutation carriers identifies five founder events and links sporadic cases to existing ALS families

**DOI:** 10.1101/685925

**Authors:** Lyndal Henden, Natalie A. Twine, Piotr Szul, Emily P. McCann, Garth A. Nicholson, Dominic B. Rowe, Matthew C. Kiernan, Denis C. Bauer, Ian P. Blair, Kelly L. Williams

## Abstract

Amyotrophic lateral sclerosis (ALS) is a neurodegenerative disorder characterised by the loss of upper and lower motor neurons resulting in paralysis and eventual death. Approximately 10% of ALS cases have a family history of disease, while the remaining cases present as apparently sporadic. Heritability studies suggest a significant genetic component to sporadic ALS, and although most sporadic cases have an unknown genetic etiology, some familial ALS mutations have also been found in sporadic cases. This suggests that some sporadic cases may be unrecognised familial cases with reduced disease penetrance. Identifying a familial basis of disease in apparently sporadic ALS cases has significant genetic counselling implications for immediate relatives. A powerful strategy to uncover a familial link is identity-by-descent (IBD) analysis which detects genomic regions that have been inherited from a common ancestor. We performed IBD analysis on 90 Australian familial ALS cases from 25 families and three sporadic ALS cases, each of whom carried one of three *SOD1* mutations (p.I114T, p.V149G and p.E101G). We identified five unique haplotypes that carry these mutations in our cohort, indicative of five founder events. This included two different haplotypes that carry *SOD1* p.I114T, where one haplotype was present in one sporadic case and 20 families, while the second haplotype was found in the remaining two sporadic cases and one family, thus linking these familial and sporadic cases. Furthermore, we linked two families that carry *SOD1* p.V149G and found that *SOD1* p.E101G arose independently in each family that carries this mutation.

## Introduction

Amyotrophic lateral sclerosis (ALS) is a severe neurodegenerative disorder characterised by the progressive loss of upper and lower motor neurons in the motor cortex, brainstem and spinal cord, resulting in paralysis and death, typically from respiratory failure, within 3-5 years of disease onset^1–5^. The majority of cases present without a family history (sporadic ALS), while 5-10% of cases are familial^6^. The cause of ALS in most cases remains unknown^7^, however heritability studies suggest a significant genetic component to sporadic ALS^8^. Furthermore, genetic mutations that are present in familial ALS cases have also been found in sporadic ALS cases^9,10^, suggesting that some sporadic cases may in fact be unrecognised familial cases with reduced disease penetrance. Identifying a familial basis of disease in apparently sporadic ALS cases has important genetic counselling implications for their immediate family members, including a 50% chance of inheriting the mutation and an increased likelihood of developing ALS.

Mutations in the gene encoding copper zinc superoxide dismutase 1 (*SOD1* [MIM:147450]) account for around 20% of familial ALS cases^2,3,5^ and a small proportion of sporadic ALS cases^9,11^. More than 150 mutations in *SOD1* have been associated with ALS thus far, where the frequency of each mutation varies across populations. The most common *SOD1* mutation in North America is p.A4V, while in Scandinavia and the United Kingdom the most common *SOD1* mutations are p.D90A and p.I114T, respectively. All three of these *SOD1* mutations, as well as *SOD1* p.D11Y and p.R115G, have originated from founder events, where the mutation has descended from a common ancestor.

Mutations that originate from founder events are typically inherited as part of larger founder haplotypes that are broken down over time due to recombination. In North America, *SOD1* p.A4V is found most often on a haplotype background that suggests it arose in American Indians. In contrast, *SOD1* p.A4V is found on a different haplotype background in Europeans, indicating two separate founder events^12^. Additionally, *SOD1* p.D90A arose from a single founder in Scandinavian families with recessive ALS, while multiple founders exist when this mutation is inherited in a dominant fashion^10,13^. Much of the work on founder events in ALS has used microsatellite markers to identify a founder haplotype^9,10,13–15^. However alternative methods are available that make use of tens-of-thousands of single nucleotide polymorphisms (SNPs) extracted from SNP array data or whole genome sequencing (WGS) data, which can also provide fine-scale resolution on the breakpoints of shared ancestral haplotypes and more accurate variant dating. These methods identify genomic regions that have been inherited from a recent common ancestor, said to be identical by descent (IBD), and have proven useful in many applications, including disease mapping^16,17^ and uncovering unknown relatedness^18,19^. In the case of founder events, individuals who have inherited part of a founder haplotype are in fact IBD over this genomic region, therefore inferred IBD regions can be used to identify common founders and thus founder events^17^.

In this study we performed an IBD analysis leveraging WGS data to investigate founder events in a cohort of 90 Australian familial ALS cases from 25 families and three sporadic ALS cases with the most common *SOD1* mutations in Australia (*SOD1* p.I114T, p.V149G, p.E101G)^20,22^. We identified multiple families and sporadic cases as distantly related and discovered several founder events in patients carrying identical *SOD1* mutations. In particular, we created relatedness networks to visualize clusters of individuals sharing a common haplotype over *SOD1*, from which we subsequently inferred the number of unique haplotype backgrounds that carry each causal *SOD1* mutation in our population, thus drawing conclusions as to the presence of founder events. This suggested that *SOD1* p.I114T and p.E101G each had two independent origins in this cohort, and p.V149G had a single origin; totalling five independent founder events. Furthermore, we were able to calculate the time to the most recent common ancestors for both p.I114T and p.V149G as less than 360 years ago.

## Material and Methods

### Australian sample cohort

850 Australian participants were recruited for analysis from the Macquarie University Neurodegenerative Disease Biobank, Molecular Medicine Laboratory (Concord Hospital), Australian MND DNA Bank (Royal Prince Alfred Hospital) and Brain and Mind Centre (University of Sydney). Each participant provided informed written consent as approved by the human research ethics committees of the Sydney South West Area Health Service, Macquarie University, or University of Sydney. Most participants were of European descent, and each ALS case was clinically diagnosed according to El Escorial criteria^21^. Genomic DNA extraction was performed from whole blood according to standard protocols. Of these 850 individuals, 90 familial ALS cases from 25 families were previously known to carry either a *SOD1* p.I114T, p.V149G or p.E101G mutation^20^. Mutation screening of the 616 sporadic cases among the 760 remaining cases determined that three sporadic ALS cases have a *SOD1* p.I114T mutation^22^.

### Whole genome sequencing data processing

Detailed descriptions of the DNA library preparation and the generation of WGS data for all 850 participants is described in McCann *et al.*^22^, as is the pipeline for processing, filtering and variant calling of this sequencing data. All 850 samples were leveraged to improve variant calling accuracy and quality filtering, however only *SOD1* mutation carriers were included for all subsequent analyses. This resulted in 88 *SOD1* samples (Table 1) and 3,527,233 high quality SNPs remaining for analysis.

**Table 1.** Familial and sporadic ALS *SOD1* mutation carrier samples.

| **Familial or Sporadic** | **Family or Sporadic ID** | **Number of Samples** | **Number of Pairs<sup>a</sup>** | ***SOD1* Mutation** |
| --- | --- | --- | --- | --- |
| Sporadic | SALS | 3 | . | p.I114T |
| Familial | 3 | 6 | 15 | p.E101G |
| Familial | 12 | 17 | 136 | p.I114T |
| Familial | 18 | 32<sup>b</sup> | 496 | p.V149G |
| Familial | 35 | 1 | 0 | p.V149G |
| Familial | 43 | 2 | 1 | p.I114T |
| Familial | 76 | 1 | 0 | p.I114T |
| Familial | 95 | 2 | 1 | p.I114T |
| Familial | 98 | 1 | 0 | p.I114T |
| Familial | 106 | 1 | 0 | p.I114T |
| Familial | 123 | 1 | 0 | p.I114T |
| Familial | 124 | 2 | 1 | p.I114T |
| Familial | 130 | 1 | 0 | p.I114T |
| Familial | 131 | 1 | 0 | p.I114T |
| Familial | 132 | 3 | 3 | p.E101G |
| Familial | 197 | 2 | 1 | p.I114T |
| Familial | 213 | 1 | 0 | p.I114T |
| Familial | 259 | 1 | 0 | p.I114T |
| Familial | 267 | 1 | 0 | p.I114T |
| Familial | 270 | 1 | 0 | p.I114T |
| Familial | 274 | 1 | 0 | p.I114T |
| Familial | 286 | 1 | 0 | p.I114T |
| Familial | 334 | 2 | 1 | p.I114T |
| Familial | 374 | 2 | 1 | p.I114T |
| Familial | mq10 | 1 | 0 | p.I114T |
| Familial | mq36 | 1 | 0 | p.I114T |
| **Total** | . | **88** | **656** | . |

<sup>a</sup>The number of pairwise comparisons was calculated for familial samples only and was simply the number of unordered 2-sample combinations, i.e. $n$ -choose-2 where $n$ was the number of samples.
<sup>b</sup>WGS data from five additional samples did not pass WGS processing quality thresholds and were not used in subsequent analyses.

### IBD analysis

Relationship estimates and IBD segments were inferred using TRIBES^23^ with default parameter settings. Briefly, TRIBES phases biallelic SNP data using BEAGLE v4.1^24^ then infers IBD segments with the phased haplotype data using GERMLINE^25^. GERMLINE identifies IBD segments by sliding a window of a predefined length along a chromosome and classifying pairs of samples as IBD within each window if they have an identical haplotype. Neighbouring windows that are inferred IBD for a pair of samples are then merged to define the IBD segment boundaries. IBD segments that overlapped the masked regions reported in TRIBES were either removed from further analyses or had their boundaries adjusted. These masked regions most likely reflect population substructure due to linkage disequilibrium and loci that are difficult to map such as centromeres^26^. We note that *SOD1* was more than 12 Mbp from its nearest masked region. IBD segments of 3cM or larger (*n*=16,736) were retained for analysis genome wide.

### Relatedness networks of shared haplotypes over the *SOD1* locus

A relatedness network is a graphical representation of shared haplotypes between pairs of individuals over a specified locus. Each node in the network represents a unique individual and an edge is drawn between two nodes if the individuals share an IBD segment, either partially or completely, over the specified locus. All individuals who do not share an IBD segment over the locus with any other individual are omitted from the network. Networks are produced using the functions getIBDiclusters and plotIBDclusters in the R package isoRelate^27^, where the network layout is produced according to Fruchterman-Reingold forced-directed layout algorithm^28^. This algorithm aims to position nodes such that all edges are of similar lengths with as few edges overlapping as possible. The locus used in this study was chr21:33,031,935-33,041,243 (hg19).

### Dating *SOD1* mutations p.I114T and p.V149G

The Gamma method^29^ was used to estimate the age of *SOD1* p.I114T and p.V149G, respectively. Variant dating could not be performed on *SOD1* p.E101G as there were too few individuals of sufficiently distant relatedness for the assumptions of the methodology to hold. Briefly, the Gamma method uses the lengths of shared ancestral haplotypes that carry the mutation to estimate the time to the most recent common ancestor, which is less than or equal to the time since the mutation first arose. Ancestral haplotype lengths were simply taken as the lengths of the inferred IBD segments generated from phased data, and the time to the most recent common ancestor is reported assuming a correlated genealogy, which takes into account subsets of samples with a common ancestor earlier than the most recent common ancestor for all samples.

## Results

### Summary statistics for the *SOD1* cohort

Following filtering procedures, 88 ALS samples and 3,527,233 SNPs genome wide were retained for analysis. Of these, 85 cases had familial ALS, where 43 individuals (21 families) carry a *SOD1* p.I114T mutation, 33 individuals (two families) carry *SOD1* p.V149G, and nine individuals (two families) carry *SOD1* p.E101G (Table 1). Additionally, three sporadic ALS cases were identified with having a *SOD1* p.I114T mutation^22^. Pairwise IBD analysis was performed on the SNP data using TRIBES^23^ and a total of 1,209 IBD segments of 3cM or greater were inferred on chromosome 21 with median length 8.31cM (range: 3cM to 62.79cM).

### New relationships identified between ALS families and sporadic cases

Of the 85 familial ALS cases, 70 came from families where multiple affected individuals were sequenced and the degree of relatedness was known (Table 1). Of these known relationships, TRIBES correctly estimated 99% of relationships to within 1 degree of the true relationship for relatives up to 7^th^ degree (third cousins), while only 13% of 8^th^ degree or higher relatives were correctly estimated to within 1 degree (Figure 1).

**Figure 1.**
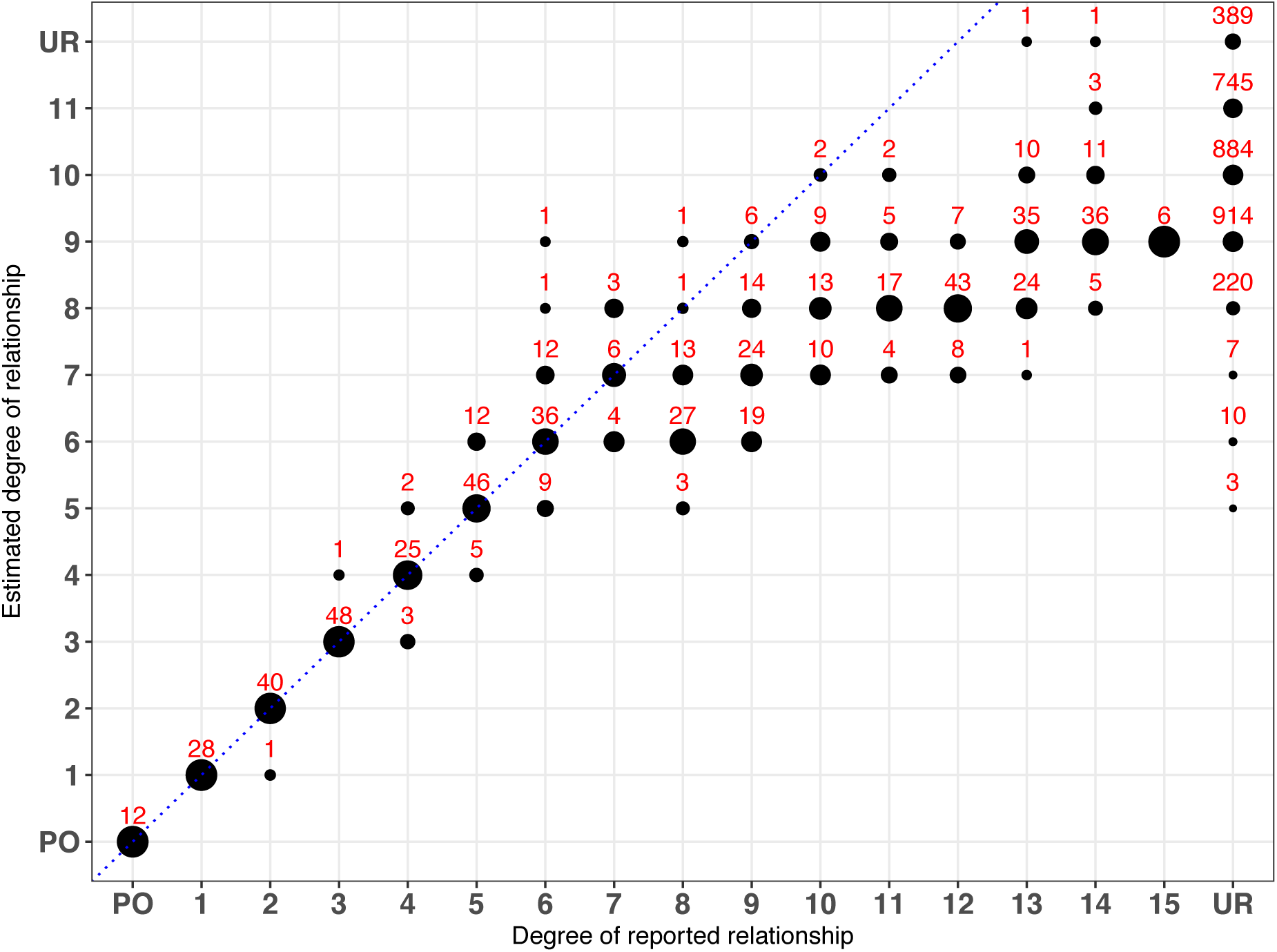
The reported vs. estimated degree of relatedness in the *SOD1* cohort using TRIBES. The size of the circles represent the percentage of individual pairs whose estimated degree of relationship are exactly the same as their reported relationship. The number of pairs estimated at each point is labelled in red above the corresponding circle. PO and UR are abbreviations for parent-offspring pairs and unrelated pairs, respectively. Individuals were reported as unrelated if they belonged to different families or were sporadic cases. Circles that fall on the blue dotted line, y=x, indicate concordance between the reported and estimated relationship. TRIBES correctly estimated 99% of relationships to within 1 degree of the reported relationships for relatives up to 7^th^ degree (third cousins) and identified 3, 10 and 7 pairs of seemingly-unrelated individuals as 5^th^, 6^th^ and 7^th^ degree relatives respectively.

By extending this analysis to identify relationships between seemingly-unrelated individuals, 3, 10 and 7 pairs of individuals were found to be 5^th^, 6^th^ and 7^th^ degree relatives respectively (Figure 1, Table 2), while there were no individuals of unknown-relatedness who were estimated as 4^th^ degree relatives or closer. Although some apparently unrelated individuals were inferred as 8^th^ to 11^th^ degree relatives (Figure 1), we chose only to investigate individuals identified as 7^th^ degree relatives or closer as this is the accuracy limit of TRIBES^23^. Of these novel relationships, 19 pairs were from patients where both individuals within each pair had identical *SOD1* variants and shared an IBD segment over this locus. This included one pair of apparently sporadic ALS cases with *SOD1* variants, which confirmed they are in fact part of a larger extended family.

**Table 2.** Newly identified 5^th^, 6^th^ and 7^th^ degree related pairs.

| **FID<sup>a</sup> 1** | **IID<sup>b</sup> 1** | **FID<sup>a</sup> 2** | **IID<sup>b</sup> 2** | **Estimated Degree** | **IID 1 Mutation** | **IID 2 Mutation** |
| --- | --- | --- | --- | --- | --- | --- |
| 18 | 18-60 | 35 | 35-142 | 5 | p.V149G | p.V149G |
| 18 | 18-58 | 35 | 35-142 | 5 | p.V149G | p.V149G |
| 197 | 197-060095 | mq36 | mq36-MQ160147 | 5 | p.I114T | p.I114T |
| 18 | 18-77 | 35 | 35-142 | 6 | p.V149G | p.V149G |
| 18 | 18-67 | 35 | 35-142 | 6 | p.V149G | p.V149G |
| 18 | 18-77 | 197 | 197-060228 | 6 | p.V149G | p.I114T |
| 334 | 334-060820 | 374 | 374-140839 | 6 | p.I114T | p.I114T |
| 334 | 334-120512 | 374 | 374-140839 | 6 | p.I114T | p.I114T |
| 334 | 334-060820 | 374 | 374-140975 | 6 | p.I114T | p.I114T |
| 334 | 334-120512 | 374 | 374-140975 | 6 | p.I114T | p.I114T |
| 123 | 123-971530 | 259 | 259-080285 | 6 | p.I114T | p.I114T |
| 197 | 197-060228 | mq36 | mq36-MQ160147 | 6 | p.I114T | p.I114T |
| SALS | MN201517 | SALS | SALS2258 | 6 | p.I114T | p.I114T |
| 76 | 76-940290 | SALS | MN201517 | 7 | p.I114T | p.I114T |
| 76 | 76-940290 | SALS | SALS2258 | 7 | p.I114T | p.I114T |
| 267 | 267-090221 | 286 | 286-090750 | 7 | p.I114T | p.I114T |
| 18 | 18-41 | 35 | 35-142 | 7 | p.V149G | p.V149G |
| 197 | 197-060095 | 43 | 43-070626 | 7 | p.I114T | p.I114T |
| 197 | 197-060095 | 43 | 43-080797 | 7 | p.I114T | p.I114T |
| 197 | 197-060228 | 43 | 43-080797 | 7 | p.I114T | p.I114T |

<sup>a</sup>Family or sporadic ID.
<sup>b</sup>Individual ID.

### Identification of five independent *SOD1* mutation founder events

Of all individuals with *SOD1* mutations, IBD segments over the *SOD1* locus were expected in 656 pairs since this was the total number of pairs known to be related prior to analysis (Table 1). However, there was more IBD sharing over *SOD1* than expected (Figure 2). We observed IBD segments in 956 pairs that indicated shared haplotypes between seemingly unrelated families and sporadic cases, where the median length of an IBD segment over *SOD1* in apparently unrelated individuals was 4cM (range: 3cM to 37.69cM).

**Figure 2.**
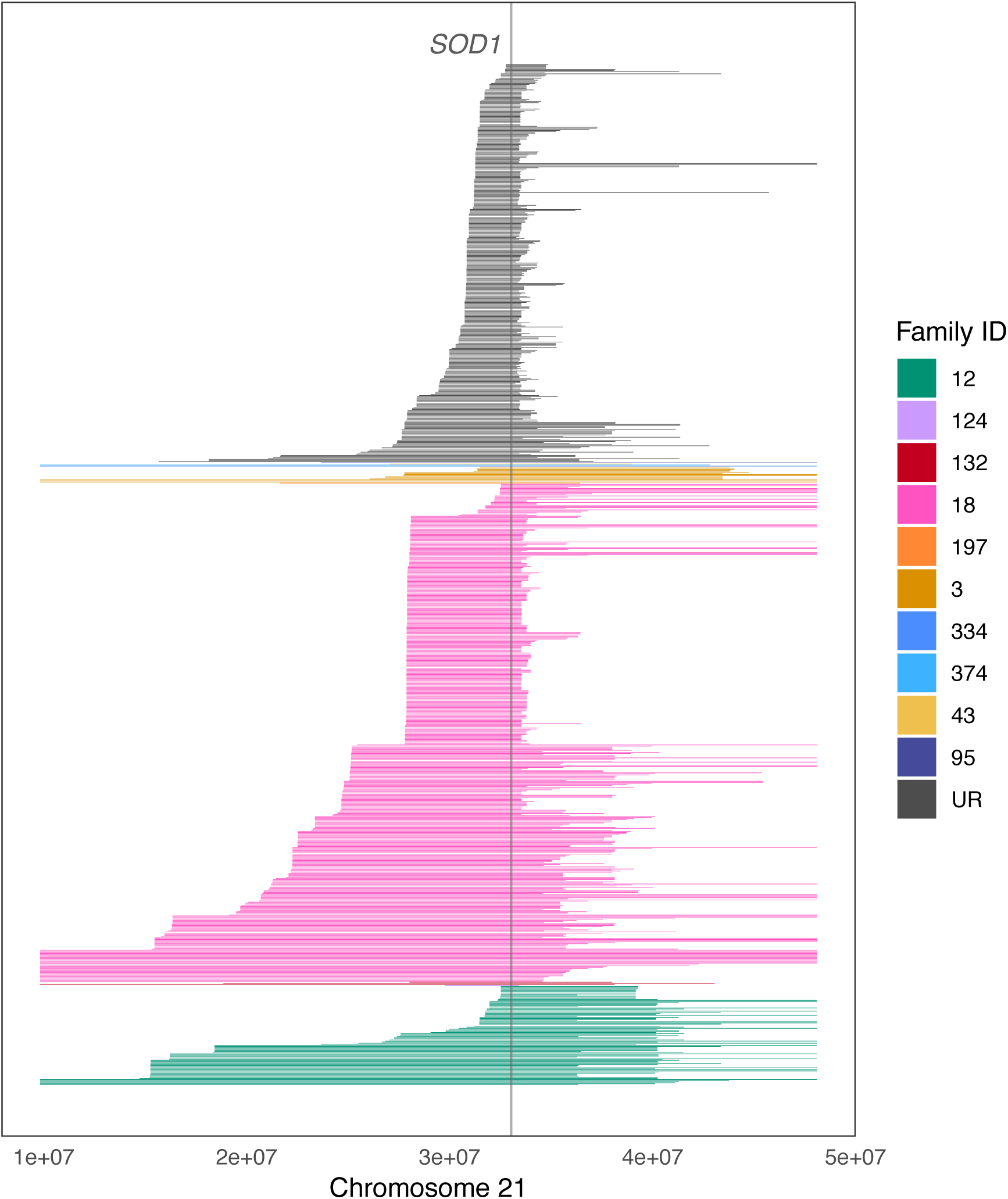
The distribution of IBD segments that overlap *SOD1*. Each line represents an IBD segment inferred between a unique pair. IBD segments have been coloured according to whether both individuals within a pair belong to the same family; or whether they belong to different families and are otherwise considered unrelated (UR). All three sporadic ALS patients with *SOD1* variants were considered unrelated. Family 18 had the greatest number of IBD segments inferred over *SOD1* as this family had the greatest number of cases sequenced, followed by family 12. Many IBD segments were inferred over *SOD1* between apparently unrelated individuals, suggesting these individuals were part of an extended family.

A relatedness network of individuals that shared IBD segments over *SOD1* is shown in Figure 3. Noticeably, five distinct clusters were evident, where every individual within each cluster carried the same *SOD1* mutation on identical haplotype backgrounds. Both families with the *SOD1* p.V149G mutation shared a common haplotype over this locus, which suggests that p.V149G descended from a common founder. Relationship estimates between cases from each family identified two pairs 5^th^ degree relatives as well as more distant relatives linking both families (Table 2, Figure 4). In contrast, *SOD1* p.E101G was found on two different haplotype backgrounds, each unique to one of the two families that carried this mutation, suggesting that p.E101G arose independently in these families. Similarly, two different haplotype backgrounds appeared to harbour the *SOD1* p.I114T mutation, implying two independent origins for this mutation in our cohort. One of these haplotypes was seen in three cases; including two apparently sporadic cases and one familial case. These three individuals were estimated to be 6^th^ and 7^th^ degree relatives. The second *SOD1* p.I114T haplotype was present in 20 apparently unrelated families as well as one apparently sporadic case, suggesting this haplotype had also descended from a common founder and was the most widely distributed haplotype in our cohort. The closest degree of relatedness estimated between families in this cluster was 5^th^ degree (Table 2).

**Figure 3.**
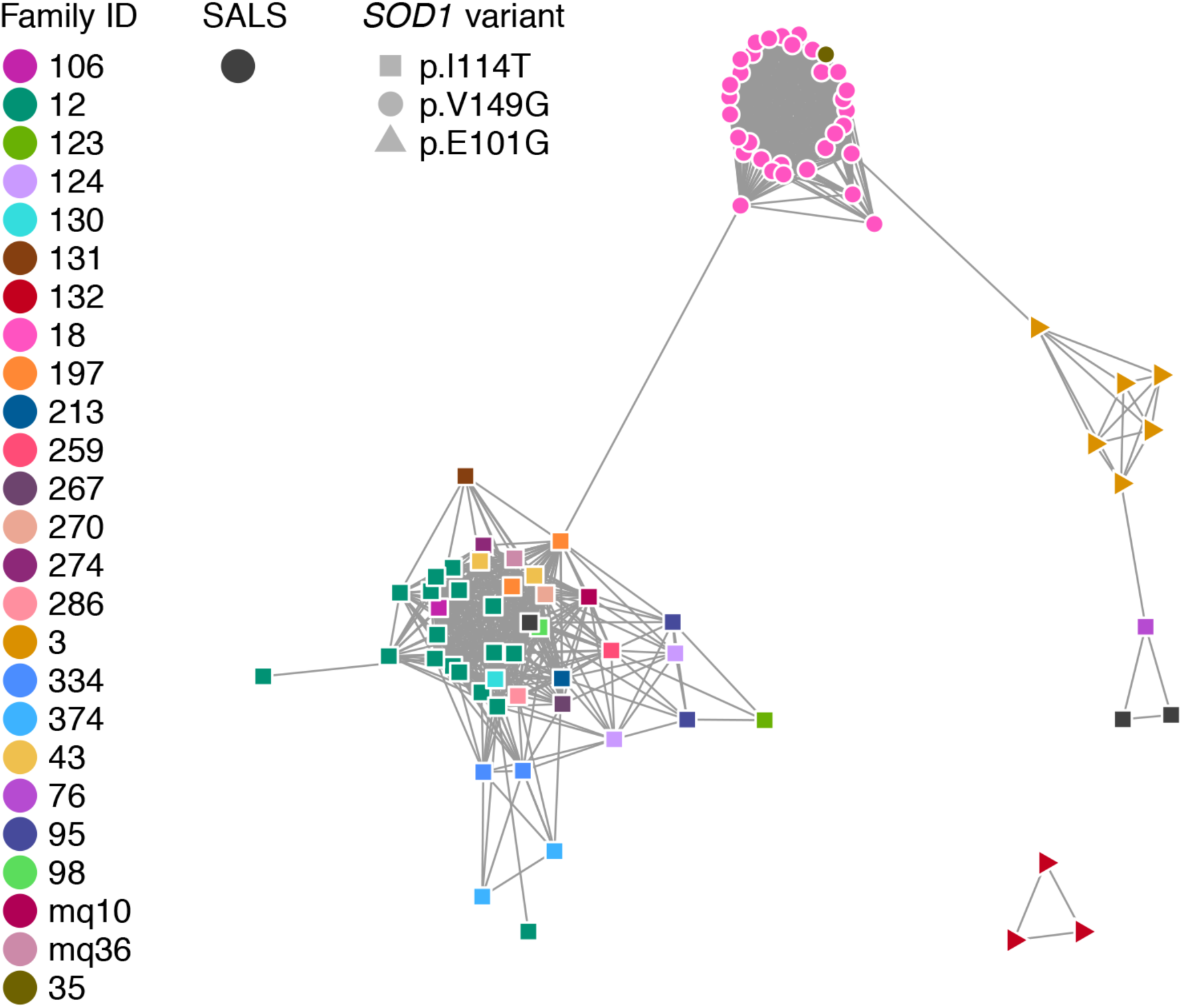
Network of individuals sharing IBD segments over *SOD1*. Each node is a sample and an edge is drawn between two samples if they were inferred IBD over *SOD1*. Nodes are coloured according to their unique family ID, in addition to the three sporadic ALS cases who have been assigned one colour. All samples have one of three *SOD1* mutations, represented by unique node shapes in the network. There are five clusters in this network, where all cases within each cluster had an identical *SOD1* mutation. The cluster of individuals carrying *SOD1* p.V149G connects family 18 and family 35, indicating they were in fact one family. Similarly, two clusters are present for individuals carrying *SOD1* p.I114T, where these individuals were from different families, including three apparently sporadic ALS cases, indicating two disjoint extended families. Specifically, two sporadic cases were found to be related to each other and family 76, while the third sporadic cases was found to be related to the remaining 20 families with *SOD1* p.I114T. In contrast, *SOD1* p.E101G was unique to each family with this mutation, suggesting independent origins. The three pairs of individuals with discordant mutations were inferred IBD over *SOD1* and likely represent false positive IBD calls.

**Figure 4.**
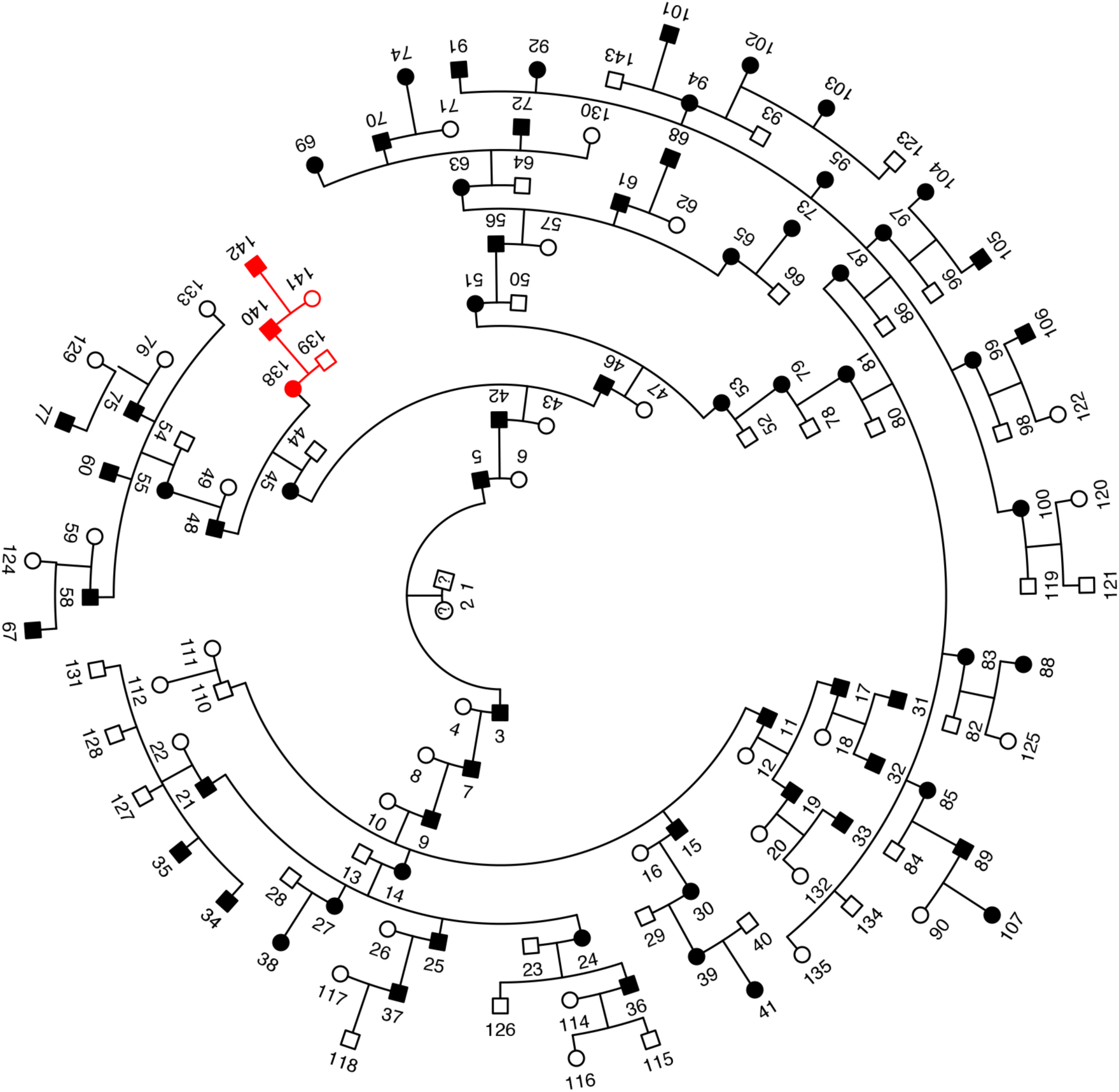
Pedigree connecting two Australian families with *SOD1* p.V149G. A subset of family 18’s pedigree (black) with 67 ALS cases over ten generations linked to family 35 (red). The extended pedigree for family 18 had 409 individuals and 67 ALS cases. The sex of individuals from generation 7 to generation 10 have been omitted for confidentiality.

### Mutation dating of *SOD1* p.V149G and p.I114T

We estimated the times to the most recent common ancestor for *SOD1* p.V149G and p.I114T, where estimation was performed separately for each of the two clusters carrying p.I114T (Figure 3). For *SOD1* p.V149G, we selected six individuals for analysis, including individuals from both families, who were at least 6^th^ degree relatives. The estimated age of p.V149G was 3 to 11 generations (60 to 220 years, assuming 20-year generation time). For the large *SOD1* p.I114T cluster (Figure 3), we selected one individual from each of the 20 families with the highest number of connections to other individuals in the network as well as the sporadic case for variant dating. The estimated age of p.I114T on the haplotype present in this cluster was between 5 to 18 generations (100 to 360 years). For the smaller *SOD1* p.I114T cluster, we included all three individuals in the calculation, and estimated the age of p.I114T on the alternative haplotype to be between 1 to 11 generations (20 to 220 years).

## Discussion

In the present study, we analyse a cohort of Australian ALS cases who have had their causal mutation, and therefore disease critical region, identified as *SOD1* p.I114T, p.V149G and p.E101G^20,22^. However as each of these three mutations appeared in multiple individuals from different families, we sought to determine if each mutation descended from one or more common ancestor. In the case of *SOD1* p.I114T, where 43 individuals from 21 families and three sporadic cases have the mutant allele, it seemed unlikely that this mutation arose independently in each family, reflecting a high mutation rate. As such, we performed an IBD analysis on WGS data to uncover any unknown-relatedness in our cohort and explore founder events.

Using TRIBES to estimate the degree of relatedness between apparently unrelated individuals, we identified 20 pairs of 5^th^, 6^th^ and 7^th^ degree relatives connecting six pairs of families, where both individuals have identical *SOD1* mutations in all but one pair. Investigating the pair with discordant mutations revealed the inferred IBD segments to be inconsistent with Mendelian inheritance (data not shown), thus they are unlikely to represent true 6^th^ degree relatives. One explanation for incorrectly identifying these individuals as close relatives is the increased number of false IBD segments produced by GERMLINE with sequencing data^30^. Many incorrectly inferred IBD segments will inflate the amount of IBD sharing observed between a pair of individuals, which in turn will give the appearance of close relatives. This may also explain why more distant relatives, such as individuals who are 12^th^ degree relatives or greater, are consistently estimated as more closely related (Figure 1).

Relatedness networks have been shown to be a powerful method to identify clusters of individuals sharing a common haplotype over a locus and can also be informative as to the number of haplotypes that segregate with disease, indicative of independent origins or founder events^17,27^. By investigating IBD segments overlapping *SOD1* using relatedness networks, we identified five distinct clusters of individuals that each carried a unique disease associated haplotype (Figure 3). Three of these clusters were each connected by one pair of individuals with discordant *SOD1* mutations, whom are unlikely to be truly related. *SOD1* p.I114T was present on two different haplotype backgrounds, one of which was inherited in 20 families and one sporadic case. p.I114T is the most common *SOD1* mutation in the United Kingdom, and in particular in Scotland^31^, where a haplotype analysis of Scottish p.I114T mutant cases revealed a common founder^9,32^. It is likely that *SOD1* p.I114T in the Australian cohort has also descended from Scottish founders, as genealogical analysis indicated that six of the p.I114T families originated from Scotland, including families in both clusters that carry different *SOD1* p.I114T haplotypes (Figure 3). Furthermore, we estimated that this mutation originated from a common ancestor up to 360 years ago, which is within the timeframe of Scottish settlers in Australia^33^.

Family 18 was the largest Australian ALS family in the cohort, spanning ten generations, 409 total individuals and 67 ALS cases with the *SOD1* p.V149G mutation^34^, of which 32 were included in this analysis. TRIBES inferred two individuals from family 18 as both 5^th^ and 6^th^ degree relatives with a single case from family 35, who also carried a *SOD1* p.V149G mutant allele. Using the relationship estimates from TRIBES along with pedigree records, we were able create a new pedigree combining both families (Figure 4). Relationship estimates combined with the inferred IBD segments confirmed that all cases with p.V149G in this cohort descended from a common founder; predicted to have originated up to 11 generations ago (220 years), which was consistent with pedigree records.

*SOD1* mutations have a large effect size^6^ and almost always present as classic ALS without comorbid frontotemporal dementia. However, the variability in disease phenotype, including age of disease presentation and duration, between individuals carrying identical mutations is marked, suggesting polygenic, epigenetic and environmental factors may also play a role in disease onset and progression. It has been postulated that separating ALS into phenotype subgroups may aid in uncovering phenotypic modifiers, whether they be genetic or epigenetic. Large ALS families with known gene mutations provide a relatively homogenous group with which to uncover modifiers. However, the late onset of ALS limits the recruitment of affected individuals, such that most recruited ALS families are represented by a small number of samples. By genetically linking families using relatedness analysis, specifically IBD sharing, we can increase family sizes and therefore increase statistical power to identify these phenotypic modifiers.

Phenotypic modifiers may also explain why some ALS cases appear as sporadic cases when they are in fact familial cases with reduced penetrance. Here, all three apparently sporadic ALS cases that carried a *SOD1* p.I114T mutation were shown to be unrecognised familial cases. This result is consistent with previous findings that familial ALS cases with *SOD1* p.I114T have been incorrectly classified as sporadic cases^9,32^. Screening these three sporadic cases for additional reported ALS causal or associated variants identified at least one other ALS mutation or associated variant in addition to the *SOD1* p.I114T mutation in each sporadic case^22^. These additional variants may be acting as disease modifiers or to reduce penetrance. In addition to incomplete penetrance, incorrect classification of sporadic ALS cases may arise from inadequate knowledge or reporting of family history and may be masked, for example, by the death of at-risk family members from other causes prior to ALS onset^6,35^. Not recognising a familial basis of disease can have significant genetic counselling implications for immediate family members^6,35^ whose risk of developing ALS greatly increases. Correct classification of familial and sporadic cases allows health professionals to make appropriate recommendations regarding genetic testing and lifestyle changes of ALS patients and their families.

Identifying relatedness and thus founder events within ALS patient cohorts aids in disease gene mapping when the causal variant is unknown. In such instances the search space for potential candidate genes can be greatly reduced to those within IBD regions common to all affected family members. Such analyses may help improve our understanding of the biological mechanisms influencing familial ALS, particularly in terms of disease progression, as well as sporadic ALS which remains largely unsolved.

## Acknowledgements

We thank Carolyn Cecere and Ashley Crook for their assistance in compiling family information. This work was funded by the Motor Neuron Disease Research Institute of Australia (grant to KLW), National Health and Medical Research Council of Australia (grant 1095215 to IPB and fellowship 1092023 to KLW) and Macquarie University (grant to KLW).

## Web Resources

The R Project for Statistical Computing, http://www.r-project.org/

R Studio, http://www.rstudio.com/

Genetic Mutation Age Estimator, https://shiny.wehi.edu.au/rafehi.h/mutation-dating/

